# Non-random mating and information theory

**DOI:** 10.1101/095901

**Authors:** A. Carvajal-Rodríguez

## Abstract

In this work, mate choice is modeled by means of the abstract concept of mutual mating propensity. This only assumes that different type of couples can have different mating success. The model is adequate for any population where mating occurs among distinct types. There is no extra assumption about particular mating scheme or preference model. The concept of mutual mating propensity permits to express the observed change in the mating phenotypes as the gain in information with respect to random mating. The obtained expression is a form of the Price equation in which the mapping between ancestral and descendant population is substituted by a mapping between random mating and non random mating population.

At the same time, this framework provides the connection between mate choice and the exact mathematical partition of the choice effects, namely sexual isolation, sexual selection and a mixed effect. The sexual selection component is the sum of the intra-sexual male and female selection.

The proposed framework helps to unveil previously hidden invariants. For example, if the mutual preference between partner types is multiplicative there is no sexual isolation (inter-sexual selection) effect on the frequencies, i.e. the only possible effect of mate choice is intra-sexual selection. On the contrary, whatever the contribution of each partner to the mutual preference, if it comes as a non-multiplicative factor, there is at least an inter-sexual selection detectable effect.

This new view over the mate choice problem, permits to develop general mating propensity models and to make predictions of the mate choice effects that may emerge from such models. This possibility opens up the way for setting a general theory of model fitting and multimodel inference for mate choice.

Thus, it is suggested that the proposed framework, by describing mate choice as the flow of information due to non-random mating, provides a new important setting for exploring different mating models and their consequences.

## 1. Introduction

Mate choice is arguably one of the most active areas of evolutionary research. There has been a lot controversy regarding the concept of mate choice. The debate around mate choice was due in part to its importance for fields so diverse as population genetics, evolutionary-ecology, animal behavior, sociology, or psychology. In addition, there has been an excess of verbal models and imprecise terminology regarding different aspects of mate choice (Edward, 2015). Mate choice can be broadly described as the effect of some expressed traits leading to non-random mating. Under this broad definition there are various aspects that can be considered. Yet Darwin (1871) distinguishes between intrasexual selection and intersexual selection. The first arises directly from competition among individuals of the same sex while the second arises from choice of mates by the other sex (Kuijper et al., 2012). Alternatively, from a population genetics point of view, mate choice is defined as the observed mating frequency deviation with respect to random mating, considering population gene or phenotype frequencies. So defined, mate choice can be partitioned into (intra)sexual selection, defined as the observed change in gene or phenotype frequencies in mated individuals with respect to population frequencies, and sexual isolation (behavioral isolation or intersexual selection), defined as the deviation from random mating in mated individuals (Rolán-Alvarez and Caballero, 2000). In this work I followed these definitions of mate choice, intrasexual and intersexual selection.

For an alternative description of these concepts and a discussion about some of the most widely used descriptions of evolutionary change within the context of sexual selection, I refer the reader to (Kuijper et al., 2012; Rosenthal, 2017).

The many aspects and complexity of mate choice justifies the extensive research that has been made in the last decades producing several theoretical models and empirical tests. Related to modeling and detection of mate choice, there is the question about the correct null hypothesis for testing the evolution of mate choice. The Lande-Kirpatrick (L-K) model has been proposed as a null model (Kirkpatrick, 1982; Lande, 1981; Prum, 2010; Roff and Fairbairn, 2014). This model assumes neutral genetic variation for the mating preference trait while the target trait can be under natural selection. However, the L-K role as a null model is not clear when the preference is set by similarity (preference and target trait coincide) and the trait is under divergent selection (Servedio et al., 2011), i.e. the trait is “magic” sensu Gavrilets (2004), because in this case the preference trait is already under selection (Hughes, 2015).

Therefore, there is still a need for both, null models and a general framework, where the key essential facts of the mate choice can be adequately described. Here, I argue that the formalism provgided by the information theory in the form of the Jeffreys’ divergence is the right tool to do so.

The information theory has already been elegantly applied for describing evolutionary change (Frank, 2009; Frank, 2012b; Frank, 2013). The present work takes advantage of that mathematical structure and applies it for modeling the change in mating frequencies due to mate choice. As far as I know there is no previous attempt of describing mate choice from the viewpoint of the information theory. Nevertheless, the potential of the informational view for evolutionary ecology has been already suggested (Dall et al., 2005).

First, I defined a general model that only requires an abstract functional relationship connecting the observed mating frequencies with the expected by random mating from the population gene or phenotype frequencies. This suffices for developing a general information equation for mate choice that can be adequately partitioned into intrasexual and intersexual information components, plus a mixed term provoked by the confounding effect of the marginal frequencies when the mating propensity effects are asymmetric. Interestingly, the three terms can be separately estimated from the observed frequencies and so, the researcher can study how different models and parameters translate into the different mate choice components. Also, it is proposed that this setting provides the baseline for solving the mate choice null hypothesis problem, since the null model emerges naturally from the idea of zero information. Thus, the correct null should not rely on neutral preference or trait genes but on zero information.

The utility of this framework is shown by analyzing a real data example. I will show how the view obtained from the unveiled relationships can be utilized to classify different general models from its consequences which facilitates the multimodel inference of the mate choice. However, a deeper study on the outcomes of different forms of the mating preference functions is out of the scope of the present article and is part of a different paper.

## 2. Model of Mate Choice

As mentioned above, the following model is as a particular specification of the information theory interpretation for evolutionary models, proposed in (Frank, 2012b; Frank, 2013). The general framework developed by this author fits perfectly for the purpose of describing the occurrence of non-random mating and the flow of information that it provokes. Remarkably, once the basic equation for the gain in information due to non-random mating is formalized, the relationship between mate choice and its different evolutionary outcomes emerges naturally, providing a clear and useful picture of the intrasexual and intersexual selection effects.

### 2.1 General model

Let consider a population with a number of *n*_1_ females and *n*_2_ males. For a given female phenotype *X*(e.g. shell color) with *K* different classes having values *X*_1_, *X*_2_ … *X*_k_, the frequency of the phenotype *X*_i_ in the female population is *p*_1i_ = *n*_1Xi_ / n_1_, i.e. the number of females with that phenotypic value divided by the total number of females. Similarly, for the male phenotype *Y* (could be the same as *X*) with *K*′ classes, the frequency of *Y*_j_ in the male population is *p*_2j_ = *n*_2*Y*j_ / *n*_2_.

In this way, by using the frequency of the phenotype for each sex, the expected mating frequencies if mating is at random is

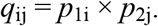

Now, given a female phenotype *X*_i_ and a male phenotype *Y*_j_, let’s define the mutual mating propensity *m*_ij_(*x, y, e*) as the number of matings of *X*_i_ with *Y*_j_ after their encounter in the environment *e*. The normalized mating propensity is

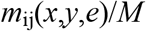

where *M* = Σ_*ij*_ *q_ij_ m_ij_* (*x,y,e*).

Then, the observed mating frequencies in a given environment *e* can be expressed as

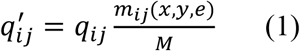

Therefore, the observed mating frequencies are the result of the functions *m*_ij_(*x,y,e*) (hereafter noted as *m*_ij_), that can be any kind of composition of the preference of female *X*_i_ for male *Y*_j_, and vice versa, in the environment *e*.

Note that random mating is a particular case of the model in (1) when the propensities are equal for every mating pair. The mutual mating propensity functions can represent empirical or analytical functions, as for example the Gaussian-like preference functions (reviewed in Carvajal-Rodriguez and Rolán-Alvarez, 2014). Moreover, each *m*_ij_ can be composed of female and male preferences, so mutual mate choice models (Bergstrom and Real, 2000) are also available under this setting. The standardized *m*_ij_ values could also be estimated a posteriori from the data. In this case they coincide with the pair total index i.e. the ratio of the frequency of the observed types divided by the expected pair types calculated from the total frequencies (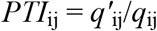, Rolán-Alvarez and Caballero, 2000) which becomes an observation of the mutual mating propensity from the mating phenotypes (see below).

Once we have the mating frequencies as defined in (1), the change with respect to random mating is

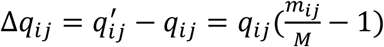

The mean population change for a combined phenotype Z = X * Y is

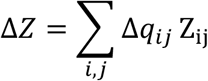

Because the relationship in (1) is defined by ratios is more natural to express the quantities in the logarithmic scale and so we can express *m*_ij_ as

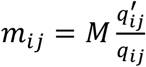

which in the logarithmic scale becomes

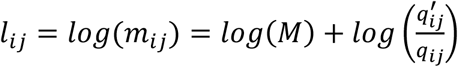

Thus, if we take the logarithm of the propensity as the combined phenotype *Z* and by noting that ΣΔ*q*_ij_ = 0 and that *log*(*M*) is constant through the summation, then we can measure the mean population change Δ*L* in relative propensity as

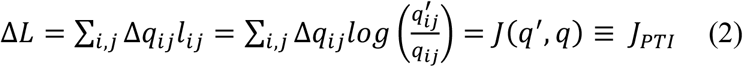

which is the Kullback-Leibler symmetrized divergence (noted as Jeffreys in Frank, 2012b), that measures the gain in information when the differential mating propensity moves the population from mating frequencies *q* to *q’* or vice versa. Note that if the propensity is equal for every pair i.e. *M* = *m*_ij_ ∀ *i,j* then *q*′ = *q* so that *J* = 0 which is the minimum information value since *J* cannot be negative.

Recall from equation (1), that each *m*_ij_/*M* is the ratio of the frequency of the observed types divided by the expected pair types from the total frequencies. This is, by definition, the pair total index *PTI* (Rolán-Alvarez and Caballero, 2000) and so, the logarithmic term in Δ*L* is the logarithm of the *PTI* values. Therefore, *J*(*q’ q*) measures the gain in information as captured by the *PTI* coefficients, confronting the hypothesis of mate choice against random mating. Hereafter, we note this *J* as J_PTI_.

Interestingly enough, the Jeffreys’ divergence computed as *J*_PTI_ (by taking the natural logarithm and multiplying (2) by the total number of matings) is well approximated by a chi-square for the null hypothesis of random mating with *KK’-1* degrees of freedom (Evren and Tuna, 2012).

The information obtained from *J*_PTI_ has been computed using the different propensities as classes for classifying the couples i.e. we considered *log(m)* as the phenotype Z. When the classes are based upon other phenotypes rather than propensities, we are conveying a specific meaning for the change in frequencies, say, the change in mating frequencies due to differential mutual propensities is observed in terms of change in shell color mating frequencies. Therefore, the phenotype can be viewed as other scale on which we can measure this information (Frank, 2013). Of course, different kinds of phenotypes can be more or less involved in mate choice and so, different scales are more or less useful for measuring the mate choice information.

### 2.2 Relative propensity and phenotypes

When we observe any mating pair (*i,j*), we need to identify the mating by a given characteristic (e.g. shell color) since we cannot directly classify it by the value of the propensity function *m*_ij_. In general, we ignore the specific form of the mutual mating propensity function *m* and so, we may assume that some phenotype matches it perfectly, as we did above (each phenotypic pair was perfectly differentiated by specific *m*_ij_ mating propensity).

Thus, if *T* is the trait that is the target of the choice, we call *J*_PTI_ to the change in the numbers of matings when these matings were classified by *T*.

Therefore, we may think on different traits *Z* that classify the mating pairs; *Z* can be a composition of female trait *X*, e.g. preference, and male target *Y*, or can be any kind of different traits or alternatively the same trait in both sexes as size, age or color. In any case, we measure the mean change in *Z* caused by differences in *m*, as

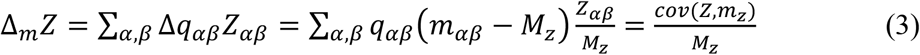

Where *cov* is the population covariance in the sense of Price (1972) as highlighted in Frank (2012a). The subscripts α, β emphasize that we are looking at pairs with observed phenotypes that not necessarily are the phenotypes exactly connected with the choice. Therefore, the propensities for the matings classified under these phenotypes can be different to the propensities for the trait *T*, then we note *m*z and *M*z to distinguish from the propensities (*m*) and mean propensity (*M*) measured directly from the real choice trait.

Equation (3) is in fact, a form of the Price equation with a different mapping for the populations involved. While the Price equation (Frank, 2012a; Price, 1972) describes the change in phenotype between two connected ancestor and descendant populations; in our equation (3), the mapping is between the random mating population and the one obtained under a given mutual mating propensity scheme.

The variable *Z* can be any desired trait including, as we assumed above, the logarithm of the propensities. So, if we take *Z* equal to the logarithm of *m*, then by substituting in (3) we obtain the mean population change Δ*L*z as

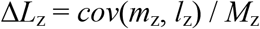

where the subscript *z* indicates that the propensities are now indexed by the trait *Z*.

Recalling the relationship in (2), we now define

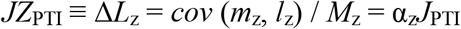

where if *J*_PTI_ > 0

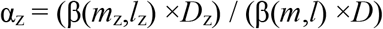

with *l*_z_ = *log*(*m*_z_), *D*_z_ = *V*(*m*_z_)/*M*_z_, *l* = *log*(*m*) and *D* = *V*(*m*) / *M* or α_z_

Note that *D* and *D*_z_ are the indexes of dispersion over the choice and *Z* traits respectively, so α_z_ is the quotient of the regressions multiplied by the index of dispersion at each phenotypic scale.

From the point of view of the estimation with real data, if we cannot measure directly the values of *m* then we simply compute *J* based on trait *Z* and therefore we are really computing *JZ*_PTI_.

In this case, note that the *PTI* coefficients are no longer the exact estimate of the mutual mating propensities because the ratio of frequencies *q′_αβ_*/*q_αβ_* does not correspond to *m*_ij_/*M* but to *m_αβ_*/*M*_z_ which is a proxy that would be more or less precise depending on the importance of the measured phenotype over the mating choice. For example, if shell size is driving mate choice, the measure of *JZ*_PTI_ (*Z* = shell size) would correspond well with *J*_PTI_ (*J*_PTI_ > 0; α_z_ ≈ 1). However, if other phenotype as shell color has nothing to do with mate choice (and is not correlated with shell size) then the measure of *JZ*_PTI_ (*Z* = shell color) would be zero (*J*_PTI_ > 0; α_z_ = 0). Further details about the distinction between *JZ*_PTI_ and *J*_PTI_ are given in appendix A.

The mate choice mediated by the differences in mutual mating propensity would produce a deviation from random mating. At the same time, this may cause two different effects, namely, intrasexual selection and intersexual selection, hereafter noted as sexual selection and sexual isolation, respectively.

### 2.3 Sexual selection

Sexual selection is defined as the observed change in gene or phenotype frequencies in mated individuals with respect to total population frequencies (Rolán-Alvarez and Caballero, 2000). This change can be studied using the frequencies within each sex, or considering jointly both sexes, by using the pair sexual selection coefficient (*PSS*, Rolán-Alvarez and Caballero, 2000). I will show that, when the *PSS* coefficients are considered a priori as the marginal propensities for the mating types, the information gained due to sexual selection is the sum of the information from each sex. When the focus is on the phenotypes instead on the propensities, the partition continue to be true, provided that the same phenotypic scale is applied when computing the *PSS* coefficients and the intrasexual components.

From the general model, the population frequency of the female phenotype *X*_i_ is *p*_1i_. The observed frequency of *X*_i_ in mated individuals, *p*′_1i_, is computed as the sum of the mating frequencies involving a female *X*_i_

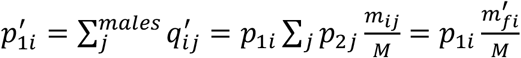

where *m*^′^_fi_ is the marginal mating propensity for the female type *i*.

Similarly for males, the frequency of phenotype *Y*_j_ is *p*_2j_ and the frequency for the male type *i* in mated individuals is

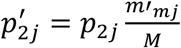

where *m*^′^_mj_ is the marginal mating propensity for the male type *j*.

The mean change in information due to sexual selection within each sex is, in terms of the female marginal propensity (female intrasexual selection)

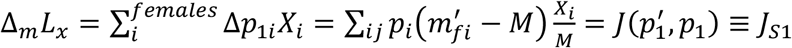

and, in terms of male marginal propensity (male intrasexual selection)

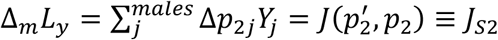

The term *J_S_* has been obtained in a similar way as for the general case, i.e. by expressing each marginal *m*^′^_fi_ and *m*^′^_mj_ in function of their respective ratio of frequencies multiplied by the mean propensity *M* and substituting the phenotype *X* or *Y*, by the logarithm of the corresponding (female or male) marginal *m*′.

The change to the scale of phenotypes produces

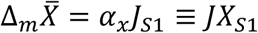

with

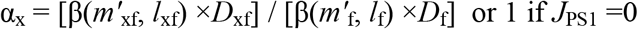

where 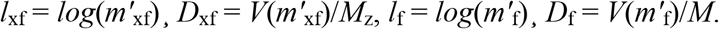.

And

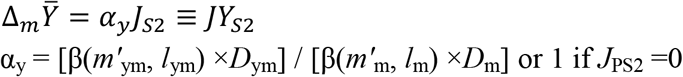

where 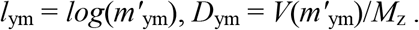.

Note that the subscripts *x* (females) or *y* (males) refer to the matings classified by phenotype instead of the true choice trait, also note that the mean of both female and male marginals is the same and equal to the mean propensity (*M*_z_ or *M* depending on the scale).

*JX*_S1_ and *JY*_S2_ are the Jeffrey’s divergence that expresses the gain of information due to intrasexual selection measured on the combined phenotypic scale *Z*.

### 2.4 Pair sexual selection

In addition to the computation within each sex, we can compare the expected pair types under random mating calculated in mated individuals, with the expected pair types from total numbers (*PSS*, see above). Thus, 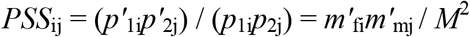. The latter term can be viewed as an a priori expression of the *PSS* coefficients. Again, the difference between the observed and the expected distribution can be expressed as

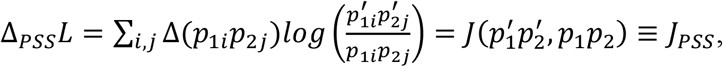

where 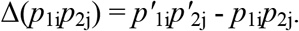.

In the scale of phenotypes

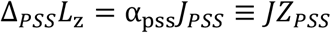

with

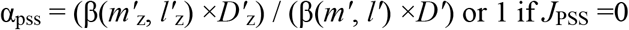

where 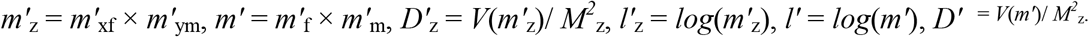.

The change in the phenotype due to sexual selection is driven by the aprioristic version of *PSS*, and is expressed in term of the information accumulated and rescaled from the marginal propensities to *Z*.

The relationship between sexual selection measured within sex and the pair sexual selection measured by *PSS* is (details in Appendix B)

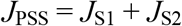

And in the scale of phenotypes

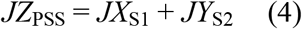

provided that the same phenotypic scale applies in the pair sexual selection statistic and in the intrasexual components (i.e. the criteria utilized for classifying the different couples is the same).

The information captured in the *PSS* coefficients is the sum of the sexual selection within each sex.

### 2.5 Sexual isolation

Sexual isolation is defined as the deviation from random mating in mated individuals (Rolán-Alvarez and Caballero, 2000). The pair sexual isolation statistic (*PSI*) is the number of observed pair types divided by the expected pair types from mates. In terms of our model this is the ratio of frequencies

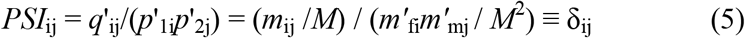

The term δ refers to an aprioristic (depends on the m’s from the model) definition of the *PSIs*. The joint isolation index for *PSI* can be expressed as

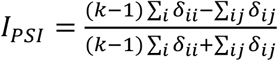

where *k* is the number of phenotypic classes involved in the classification of the matings (Carvajal-Rodriguez and Rolan-Alvarez, 2006).

As with the previous pairwise statistics, we may obtain the equations of change between observed and expected pair types in terms of *J*.

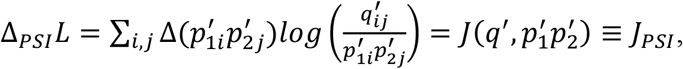

where 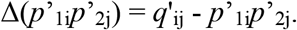.

In the scale of phenotypes

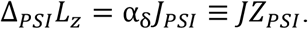

The scaling factor α_δ_ is not always easy to compute. Provided that there is no sexual selection (*J*_PSS_ = 0) then α_δ_ = α_z_ and so *JZ*_PTI_ = *JZ*_PSI_ = α_z_*J*_PSI_. Otherwise we need to rescale the factor *E*_0_ (see below) to finally get the transformation between *JZ*_PSI_ and *J*_PSI_ (see Appendix *C*).

The *JZ*_PSI_ index provides the correct metric to express the part of change in mating information that translates into sexual isolation. Presenting the *PSIs* under this formalism allow us to appreciate some facts that are not obvious from the a posteriori definition of coefficients estimated from data. We must realize (see equation 5) that if the normalized propensity of each pair (*m*_ij_ / *M*) is the product of the normalized marginal types of each partner then *δ* = 1 and so, both, the values of *I*_PSI_ and *J*_PSI_ are zero indicating no sexual isolation at all. Thus, in any model in which the mutual mating propensity is multiplicative, the only possible outcome from mate choice is intrasexual selection.

We can illustrate the multiplicative effect by means of a simple model based on a real species scenario. The bird sage grouse *(Centrocercus urophasianus*) has elaborate courtship rituals. In the spring season, males congregate in leks that are visited by the females that actively choose one of the males for mating. The number of females visiting a male seems to be related with the male long-range acoustic broadcasts whereas the probability of mating once visited is related to the visual display (Gibson, 1996).

It has been suggested that both traits, acoustic broadcast and display rate, yield a multiplicative preference for males with specific acoustic conditions and high display rates (Gibson, 1996; Rosenthal, 2017).

Thus, we can define a model of the multiplicative effect of the aforementioned traits (see details in Appendix D). Obviously, the real mating scenario is by far more complex, but the example suffices to illustrate the point.

The females are the choosers and so our model assumes a single female phenotypic class (*X*) and two male traits with two phenotypic classes each, *B/b* for acoustic broadcast, and *D/d* for display rate, where in both cases the upper case refers to the higher value of the trait. We define a multiplicative preference effect for acoustic broadcast and display rate, so that the female propensity for males *BD* can be expressed as the product of the female propensities for *Bd* and *dB*, i.e. *m*_XBD_ = *m*_XBd_ × *m*_XbD_ (supplementary Table S1).

Under this model, the mean propensity *M* coincides with the female marginal, *m*^′^_fX_ = *M*. The four male marginal propensities (*m*^′^_mBd_, *m*^′^_mBD_, *m*^′^_mbD_, *m*^′^_mbd_) have the same values as their corresponding mutual propensities (*m*_XBd_ = α, *m*_XBD_ = αβ, *m*_XbD_= β, *m*_Xbd_= 1; see Appendix D).

The model is multiplicative since each normalized mutual propensity is equal to the product of the normalized marginals, e.g. *m*_XBd_ /*M* = (*m*^′^_fX_/*M*) × (*m*^′^_mBd_ /*M*).

By computing the aprioristic expressions in (5), we see that δ_Bd_ =δ_BD_ = δ_bD_ = δ_bd_ = 1.

Thus, provided that the mating reflects the propensities, the result is that independently of the phenotypes, the number of observed pair types would be equal to the expected pair types from mates, which means that there is no sexual isolation.

On the other hand, the model predicts male sexual selection whenever *m*_XBd_ and/or *m*_XbD_ ≠ 1.

## 3. Relationship between Mate Choice, Sexual Selection and Sexual Isolation

The information as captured by the *PTI* coefficients can be partitioned in terms of *PSS* and *PSI*. Recall the expression (2) for *J*_PTI_

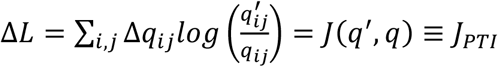

The term Δ*q*_ij_ can be expressed as the sum of the frequency changes for sexual selection and isolation

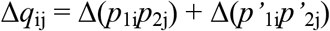

The logarithmic term 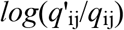 which we have also noted as *log*(*PTI*) is also partitioned in the sexual selection and isolation components

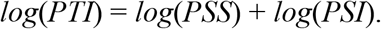

Therefore

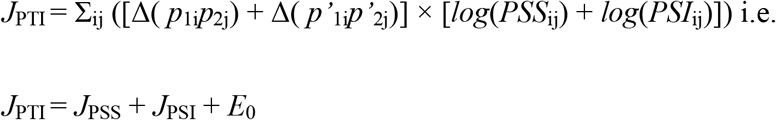

where 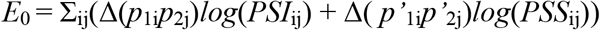. However, note that 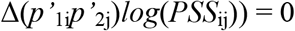 (see Appendix E) so finally

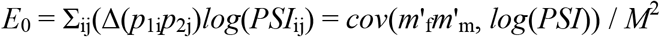

thus *E*_0_ is proportional to the covariance between the marginal propensities and the logarithm of the *PSIs*.

The covariance expression is useful for defining a scaling factor (see Appendix C) i.e.

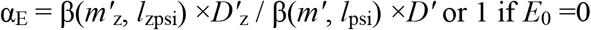

where *l*_zpsi_ = *log*(*PSI*_z_), *l*_PSI_ = *log*(*PSI*) and *m*^′^_z_, *m*^′^, *D*^′^_z_ and *D*′ are the same as defined for α_pss_. The subscript in *PSI*z indicates that this is the value obtained under trait *Z* contrary to *PSI* which is obtained directly under the choice trait.

Then, α_E_ permits to interchange between the scale of phenotypes and choice so, *ZE*_0_ = α_E_E_0_.

Alternatively, we can also express *E*_0_ (see Appendix E for details) as

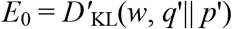

which is a Kullback–Leibler-like divergence with weights *w*_ij_ = (*PSS*_ij_ – 1)/ *PTI*_j_ in the observations *q*′. Note that contrary to the standard K-L divergence, *E*_0_ can be negative depending on the weights.

The total information is separated into the sexual selection (*J*_PSS_) and isolation (*J*_PSI_) components plus the mixed term *E*_0_. Note that *E*_0_ appears only when both sexual selection and sexual isolation effects occur so that the above given covariance is not null.

If *E*_0_ =0 this means that *J*_PSS_ and/or *J*_PSI_ capture the complete information from mate choice. When *E*_0_ is positive it indicates that the information gathered from *J*_PSS_ and *J*_PSI_ separately is not the total information from mating choice. On the other side, when *E*_0_ is negative there is some inverse relationship between sexual selection and sexual isolation information.

In the scale of phenotypes the partition still holds provided that the same phenotypic classification is applied when computing the different measures

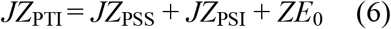

where *ZE*_0_ is the value of *E*_0_ in the given phenotypic scale.

For any given logarithmic base, the amount of the total information, *JZ*_PTI_, depends on the magnitude of the differences among the mutual mating propensity values linked to the population phenotypes under study. The higher the differences encountered the higher the value of *JZ*_PTI_. Without loss of generality, from herein we consider the natural logarithm because this facilitates testing against the null hypothesis of no information by means of the chi-square distribution.

We have given formulae for the change in the phenotypic scale for every term in (6) except *JZ*_PSI_. In this case, we have to predict the change in the scale by computing the remaining factors and solving for *JZ*_PSI_ (Appendix C).

If, as expected, the observations used to compute the information statistics come from the same sample, the sum in (6) is exact so it recovers the whole information gathered from mate choice. On the contrary, if the computations has been performed using different samples, it could be a remaining part of mate choice information that is non-explained by the above statistics but that can be recovered by the error term

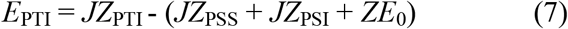

that reflects how much information may be lost due to differences in the measurement of the involved phenotypes when computing the different information components from separate samples.

## 4. Real Data Application

The theoretical framework I have presented so far has been defined in a general way, for any number of male and female phenotypic classes, and for any kind of mutual mating propensities. The application of the *J* statistics to a data sample of dimorphic traits (two classes), is immediate. For clarity, I will use the same example that appears in the pairwise statistics (*PTI, PSI* and *PSS*) original article (Rolán-Alvarez and Caballero, 2000). The correspondence between the pairwise statistics notation used by (Rolán-Alvarez and Caballero, 2000) and ours is as follows. The two phenotypic types are noted as *A* and B, the total number of observed matings is *t* and the number of *A* type females (A′ in Rolán-Alvarez and Caballero, 2000) becomes, under our notation, *p*_1A_*n*_1_, and so *B* is *p*_1B_*n*_1_; the number of *A* males becomes *p*_2A_*n*_2_ and *B’* males are *p*_2B_*n*_2_. The observed absolute number for each pair (*i, j*) would be *q*^′^_ij_*t* with *ij* ∈ {*A, B*} (see Table 1). The total number of expected mating pairs from population frequencies is *n*_1_*n*_2_ corresponding to the quantity *S* in (Rolán-Alvarez and Caballero, 2000).

**TABLE 1.**
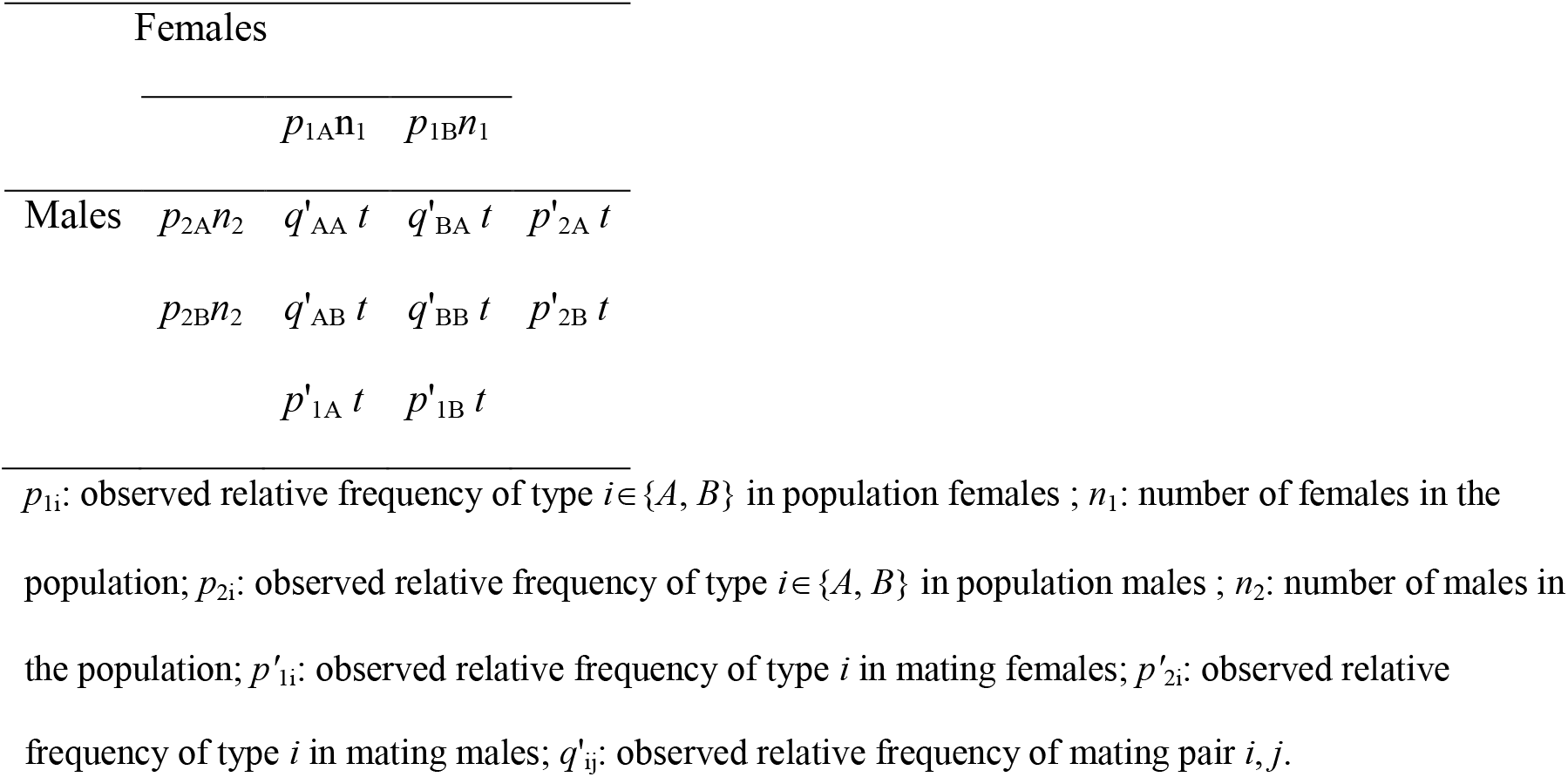
The mating model for two phenotypic classes identified as types *A* and *B*. The number of observed mating pairs is *t*.

The data correspond to a multiple-choice experiment involving two different lines of *Drosophila melanogaster* so calledM-like and Z-like (Hollocher et al., 1997). Rolán-Alvarez & Caballero applied the pairwise statistics to this data and confirmed the previous results from Hollocher *et al* indicating stronger sexual isolation than sexual selection. They also suggested a fitness advantage of females versus males but they were not able of distinguishing between female sexual selection and male preference for *M* females.

To perform the analysis, we expressed the observed data from that experiment in terms of the information model as presented in Table 1. In doing so, and noting that the observed number of mating pairs was *t* = 1704, we obtained the necessary quantities in terms of our model (Table 2).

**TABLE 2.** Analysis using the mate choice information model (Table 1 and equations 7) on *D. melanogaster* mating data from (Hollocher et al., 1997). The number of observed copulating pairs is *t* = 1704.

|  |  | Females Z | Females M |  |
| --- | --- | --- | --- | --- |
| | | $0.5 \times 1440$ | $0.5 \times 1440$ | |
| Males Z | $0.5 \times 1440$ | $0.3585 \times 1704$ | $0.145 \times 1704$ | $0.5035 \times 1704$ |
| Males M | $0.5 \times 1440$ | $0.051 \times 1704$ | $0.4455 \times 1704$ | $0.4965 \times 1704$ |
| | | $0.4095 \times 1704$ | $0.5905 \times 1704$ | |
| $JZ_{PTI}$ | 0.526 | | | |
| $JZ_{PSI} (I_{PSI})$ | 0.468 (0.63) | | | |
| $JX_{S1} + JY_{S2}$ | $0.033 + 0.00005$ | | | |
| $ZE_0$ | 0.024 | | | |
| $E_{PTI}$ | 0 | | | |

The total mate choice information obtained in *JZ*_PTI_ is partitioned in 89% of sexual isolation (*JZ*_PSI_ / *JZ*_PTI_ = 0.468 / 0.526 = 0.89; *I*_PSI_ = 0.63), 6% of sexual selection and 5% of mixed effects which explains the 100% of *JZ*_PTI_. The information coming from sexual isolation is 14 times that from sexual selection, result that matches pretty well the outcome in (Rolán-Alvarez and Caballero, 2000).

The value of *JZ*_PTI_ multiplied by the number of matings can be approximated by a chisquare variable with 3 degrees of freedom under the expectation of *JZ*_PTI_ = 0, the *p*-value obtained was below 0.00001 which indicates non-random mating. The test against *JZ*_PSI_ =0 with 1 degree of freedom, also had a p-value below 0.00001. The test against *JZ*_PSS_ =0 was also below 0.0001. However, testing separately the female and male sexual selection cases (with one degree of freedom each) produced a p-value below 0.0001 for the female case but a p-value of 0.77 for males.

Thus, we detected significant sexual isolation and selection effects as previously reported by (Rolán-Alvarez and Caballero, 2000). The sexual selection component is caused by a significant intrasexual effect in females. The mixed term *E*_0_ is positive thus indicating that not all the information is recovered by the *PSS* and *PSI* coefficients. This is due to the confounding effect which explains as far as the 5% from the total information.

### 4.1 Exploring models

In the analysis performed above we used the information partition for testing if the observations can be explained by random mating, in a similar way as we do when using the I_PSI_ statistic for testing sexual isolation (Carvajal-Rodriguez and Rolan-Alvarez, 2006; Rolán-Alvarez and Caballero, 2000).

However, the proposed theoretical framework permits going further than just testing random mating. We can rely on the described properties of mutual propensities under sexual selection and isolation, for defining different effects models. If we can define models from which we can predict the effects, then we can try to fit and infer significant parameters from the available data.

As an example, I have used the software InfoMating (Carvajal-Rodriguez, 2017) to estimate the mutual mating propensity parameters associated to the data in Table 2. The software uses the *J* information framework to a priori construct (before data) different effects models, and then compare the fitting of random mating, sexual selection and sexual isolation models to the data. There are models having sexual selection only in females, only in males or in both. The models with sexual isolation will have or not sexual selection depending on the frequencies (the conditions on marginal propensities for sexual selection are frequency dependent). The most complex model is also considered. Under this model the mutual mating propensities are estimated by the *PTIs* that are indeed the maximum likelihood estimates.

I have considered BIC (Schwarz, 1978) and AIC (Akaike, 1973) selection criteria. Both gave similar results. The best fit model was a two parameter model with sexual isolation and female sexual selection effects. The model uses two parameters *a* and *b* to define the four mating propensity values as *m*_ZZ_ = *a*, *m*_ZM_ = 1-*b*, *m*_MZ_ = 1, *m*_MM_ = *a* + *b*.

The obtained estimates under the BIC criterion were *a* = 2.47 and *b* = 0.64 which after normalization, provide the mutual mating propensity estimates as they appear in Table 3.

**TABLE 3.**
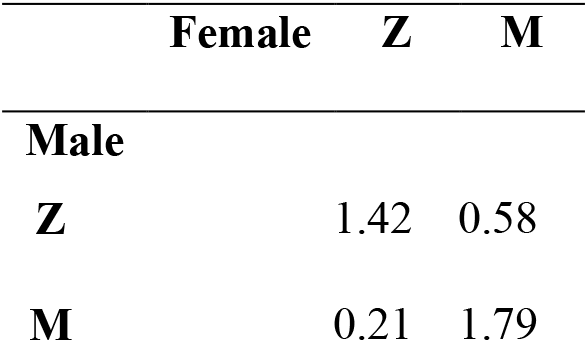
Mutual-propensity estimates from multimodel inference.

The obtained estimates are almost identical to the corresponding *PTI* values but we have only needed two parameters instead of three for defining the model. Therefore, the two parameter model may provide some insight into the biology of the mating relationships.

The obtained estimates imply positive assortative mating because the homotype mutual propensities (main diagonal in Table 3) are higher than the heterotype ones (antidiagonal, *m*_ZM_ and *m*_MZ_). If we compare the mean homotype versus the mean heterotype mating propensities, the difference is *a* + *b* -1. The value 1 is the value under random mating so, the increase of homotype mating with respect to random mating is *a* + *b*.

Moreover, the chosen model has no male sexual selection effect. This is clear when we measure the mean effect of changing the male type in the matings. We see that the effect is 0 i.e. (*m*_MM_ – *m*_MZ_ + *m*_ZM_ – *m*_zz_) / 2 = 0. On the contrary, the mean effect of changing female Z by M is *b*.

Thus, the deviation from random mating in the data from Table 2 is composed of a sexual isolation effect captured by the parameter *a* plus an effect *b* of sexual selection focused on the M females which may imply that this females are more receptive to mating in general.

## 5. Female preference and male display models

The example we have considered involves the same trait in female and male. However, there are several situations where the female preference is for a male display trait (Pomiankowski and Iwasa, 1998). In this case, the female trait is the exerted preference and the male trait is the target phenotype. In the preference-display context, the traits involved are different between sexes so that the crosses cannot be classified in homotypic versus heterotypic, which prevents the calculation of I_PSI_ and other similar indices that are only applicable to mating models in which the female and male phenotype is the same (similarity/dissimilarity models).

The mutual mating propensity framework can easily capture the preference-display scenario to express the components of mate choice in terms of information.

In Table 4 we appreciate three examples of such preference-display models. There are two types of females which have preference for males displaying phenotypic values *A*, *B* or *C*. The frequencies for the different phenotypes are equal. The mating propensities have been defined with only one parameter and three possible values, namely *a, a*/2 or virtually 0 (ε). In the first column the female preference generates a situation of complete isolation; in the second column the resultant effect of the female preference is of full intrasexual selection in males and the third column corresponds to a mixed scenario were both sexual selection and isolation occur with a mixed effect of -24% than indicates an strong overlap between both effects.

**TABLE 4.** Mating propensity models of female preference for male display traits. Two types of females ‘0’ or ‘1’ might have different preferences for males presenting distinct values for some secondary trait (*a* = 1, *a*/2 or ε). Females are the choosy sex so that the generic model implies only the female acceptance (or preference) function *f*_1_.

| $m_{\text{female},\text{male}}$ | Isolation | Sexual selection | Mixed | Generic |
| --- | --- | --- | --- | --- |
| $m_{0A}$ | $a$ | $a$ | $a$ | $f_1(0,A)$ |
| $m_{0B}$ | $a/2$ | $a/2$ | $\epsilon$ | $f_1(0,B)$ |
| $m_{0C}$ | $\epsilon$ | $\epsilon$ | $\epsilon$ | $f_1(0,C)$ |
| $m_{1A}$ | $\epsilon$ | $a$ | $\epsilon$ | $f_1(1,A)$ |
| $m_{1B}$ | $a/2$ | $a/2$ | $\epsilon$ | $f_1(1,B)$ |
| $m_{1C}$ | $a$ | $\epsilon$ | $a$ | $f_1(1,C)$ |
| $JZ_{\text{PTI}}$ | 13 | 13 | 26 | |
| $JZ_{\text{PSI}}/JZ_{\text{PTI}}$ | 100% | 0% | 75% | |
| $(JX_{\text{S1}}+JY_{\text{S2}})/JZ_{\text{PTI}}$ | 0% | 100% (0% + 100%) | 49% (0% + 100%) | |
| $ZE_0/JZ_{\text{PTI}}$ | 0% | 0% | -24% | |

$\epsilon$ : represents an infinitesimal value for which the quantity $\epsilon\log(\epsilon)$ will be zero.

However, upon inspecting the propensities in the Table 4, the effects (isolation, selection, and mixed) of the preference-display scenarios are not so intuitive, which stresses the usefulness of the information partition. For example, the column “Mixed” can be represented in a two-way table (Table 5).

**TABLE 5.** Mating preferences involved in the mixed model from Table 4.

| Males | A | B | C |
| --- | --- | --- | --- |

| Females |  |  |  |
| --- | --- | --- | --- |
| 0 | a | $\varepsilon$ | $\varepsilon$ |
| 1 | $\varepsilon$ | $\varepsilon$ | a |

The pattern in Table 5 is a clear case of isolation that splits females 0 and males A from females 1 and males C. Recall that a mixed model implies isolation+ sexual selection. The model is mixed because there is strong sexual selection against B males that virtually do not mate. At this stage, we do not care if this is because A and C have more vigour than B in the searching for mates, or because females in general do not like B males. The result is male sexual selection (against B males), so the model is mixed because the preferences in the model produce both sexual isolation and selection.

We can perform a similar exercise with the other models in Table 4 and see, for example, that the isolation model provokes isolation because females 0 prefer A while skip C, and vice versa, females 1 skip A and prefer C (both female phenotypes have the same preference for B).

Under uniform frequencies in both sexes, this isolation model does not generate sexual selection. The marginal propensity of females 0 and 1 is the same, *m*_′f0_ = *m*_′f1_ = (*a* + *a*/2 + ε) / 3 so there is no sexual selection in females. In males, the marginal propensity of A and C is (*a*+ *ε*)/2 while is *a*/2 for B males; they are equal except for the addition of the factor ε/2 which is virtually 0 and therefore there is no detectable effect of sexual selection.

## Discussion

The mate choice model defined in (1) is valid for phenotypes and genotypes, and it only requires the abstract representation of any kind of relative mutual mating propensity.

The model in (1) is similar to the model for the mating pattern predicted from encounter-mating (EM) scenarios when the availability of individuals is not affected by the matings that have already occurred (equation 19 in Gimelfarb, 1988). The latter happens in polygamous species, or even with monogamous, when only a small fraction of individuals of both sexes successfully mate (i.e. the process of the encounter corresponds to sampling with replacement).

On the contrary, when the species are monogamous and the population size is small, the mating pattern will depend on the kind of pair formation process (Gimelfarb, 1988). In the latter case, the information framework should still be valid but the equations must be updated after each mating round. Therefore, the pair formation process without replacement, would introduce some noise in the obtained mating patterns. The application of the proposed methodology in such situations is left for future work.

At the same time, (1) is analogous to the Wright’s selection equation for the change in gene frequencies so, from the viewpoint of that analogy, the relative propensity would play the role of fitness referred to each mating couple. By defining the relationship between observed and expected mating frequencies as a function of relative mating propensity, the choice is expressed as a potentiality which is also a key characteristic of fitness (Wagner, 2010).

As with the fitness concept, the mate propensity faces two main aspects, namely the measurement of differences between couples, and the intrinsic causes that provokes the propensity values. By expressing the equation of change in terms of the choice information and its components, this work focused in the first aspect.

I have connected the cause of mating choice, which is modeled by the abstract concept of mutual mating propensity, with the different possible outcomes. Notably, the connection between mate choice and its consequences appears in terms of information. The general equation (*J*_PTI_) represents the information gained by mate choice with respect to random mating. This general information is the sum of the information due to sexual isolation and sexual selection, plus a mixed effect term that can be computed separately from the others. The mixed term measures the adjustment of the partition components with respect to the total mate choice information. In addition, the information from sexual selection is the sum of the male and female intrasexual selection information.

Although the model has been constructed assuming discrete phenotypes, it is possible to estimate the Kullback-Leiblerg divergence for the continuous case (Pérez-Cruz, 2008) in order to apply a similar mate choice information partition for quantitative traits.

The information framework also provides a baseline for defining adequate null hypotheses for the distinct aspects of the mate choice problem. In fact, the information terms are mean log-likelihood ratios, so we can apply them for contrasting the different null hypothesis about random mating, sexual selection, and isolation.

Therefore, the statistical test defined as *nJ* (total number of matings *n*, multiplied by the Jeffreys’ divergence) is similar to a G-test. In fact, if we note *G* for the *G*-test with *nq* expected counts, and *G*′ for the *G*-test with *nq*′ expected counts, then *nJ* = (*G* + *G*′) / 2.

Indeed, it has been shown that the G-test G, is highly correlated (0.99) with the Jeffreys’ statistic (Evren and Tuna, 2012).

We can perform the test against random mating by considering a chi-square distribution with *KK*′-1 degrees of freedom (Evren and Tuna, 2012; Sokal and Rohlf, 1981), where *K*×*K* is the number of different mating categories. The intrasexual selection components correspond to *K*-1 and *K*′-1 degrees of freedom for *K* female and *K*′ male traits respectively. In addition, the sexual isolation component corresponds to (*K*-1)(*K*′-1) degrees of freedom.

Of course, we may also use randomization tests if we prefer to rely on the empirical distribution approach.

Therefore, if we want to contrast mate choice for a given trait *Z*, we test deviations from zero information in *JZ*_PTI_ and its components. However, if we want to contrast mate choice in general, we must test deviations from zero information in *J*_PTI_ which should be the same that testing a flat preference function across all trait values (Edward, 2015).

In addition to contrasting the null hypothesis of random mating, we may take advantage of the informational partition of mate choice effects to develop different kind of general models defined by their effects. This is possible because the developed relationships expose and clarify useful general properties, such as the requirement of non-multiplicative mutual propensity functions for obtaining sexual isolation effects and the connection of the marginal propensities at each sex with sexual selection.

As an example of the possible insight that can be gained relying in the informational framework, I reanalyzed the well-known example of D. *melanogaster* mating data from (Hollocher et al., 1997) and besides confirming previous results on the components of sexual isolation and selection effects, I have been able to fit a simple two-parameter model that explains the data by means of a component of sexual isolation plus a sexual selection component favoring the mating of the M-type females.

In addition to the similarity models in which the same phenotype is involved in both sexes, the preference-display models are also easily interpreted in terms of information and we have been able of inspecting models of full isolation, full intrasexual selection, and mixed effect models.

We have also seen an example with multiplicative mutual propensity by means of a simple preference-display model based on the bird sage grouse *(Centrocercus urophasianus)* in which the traits ‘acoustic broadcast’ and ‘display rate’, act multiplicatively over the preference.

To conclude, it is worth mentioning that the concept of mate choice is important in the evolutionary theory and other disciplines. It has been approached from a diversity of fields and inference methodologies, which has provoked that the terminology has not always been very precise. This may have contributed to some confusion in terms of causes and effects jointly with plenty discussion (Ah-King and Gowaty, 2016; Edward, 2015; Janicke et al., 2016; Roughgarden et al., 2015).

Here, I have shown that the mean change in the mating phenotypes can be expressed as the information gained due to mate choice. Overall, the obtained results lead to the suggestion that the information interpretation of mate choice is an interesting avenue that may help to improve the study of the causes as well as the effects of this important evolutionary phenomenon.

## Acknowledgement

I would like to thank Carlos Canchaya and two anonymous reviewers for their valuable comments on the manuscript. This work was supported by Xunta de Galicia (Grupo de Referencia Competitiva, ED431C2016-037), Ministerio de Economía y Competitividad (BFU2013-44635-P and CGL2016-75482-P) and by Fondos FEDER (“Unha maneira de facer Europa”).

## Appendix A. Relative propensity and phenotypes

Let *T* the trait that is the target of the choice while *Z* is any other trait that can be more or less related to the choice.

We call *J*_PTI_ to the change in the number of matings when these matings were classified by *T* (equation 2 in the main text). On the other hand, the change in the number of matings when they were classified by *Z* is *JZ*_PTI_.

If we cannot measure directly the values of *m* (trait *T*) then we simply compute *J* based on trait *Z* and therefore we are really computing

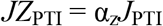

The scaling factor α_z_ is

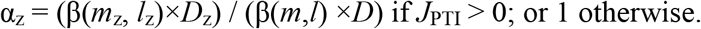

The values *D*_m_ = *V*(*m*) / *M* and *D*_z_ = *V*(*m*_z_)/*M*_z_ are the indexes of dispersion over *T* and *Z* respectively; the subscript *z* indicates that the matings were classified by phenotype Z instead of by *T* (see Appendix C for more details in the scaling formulae).

The distinction between *JZ*_PTI_ and *J*_PTI_ matters because when the information produced by mate choice is computed as *JZ*_PTI_, a value of zero could means that *i*) *cov*(*m*_z_, *l*_z_) = 0 so α_z_ = 0 i.e. the trait *Z* do not covariate with the differential propensities (the mating is random with regard to *Z*) or, alternatively ii) *J*_PTI_ = 0 meaning that there is no differential mating propensity at all, i.e. the mating is random independently of the trait we focused on.

Let’s see an example of the first situation i.e. there is mate choice but the trait *Z* is not involved in the mate choice process. Thus, assume that some unknown trait *X* that is involved in an assortative mating process exists. There are two phenotypic classes ‘1’ and ‘2’ so that *m*_11_ = 2, *m*_12_ = 1, *m*_12_ = 1, *m*_22_ = 2; the phenotype frequencies are uniform in males and females, *p* = 0.5, and mean propensity *M* = 1.5. This results in *J*_PII_ = *J*_PSI_ = 0.1155.

However, when counting the matings, we evaluated a phenotype *Z* with classes *A/B* that are independent of the mating choice process. If the trait responsible of the mate choice is uniformly distributed over the phenotypes *A/B* (i.e. half of *A* individuals have trait value ‘1’ and the other half have value ‘2’ and the same is true for *B* individuals) then the expected preference for the phenotype pairs are

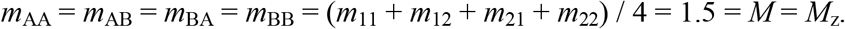

Thus it is clear that the normalized preferences are 1 and *JZ*_PTI_ =0.

Consider now a different case in which *JZ*_PTI_ ≠ 0, this means that the non-random mating is correlated to some extent with the trait *Z*. For example, consider the same mate choice scenario as above with *J*_PTI_ = *J*_PSI_ = 0.1155 but now the phenotype under study is partially linked to the mate choice so *m*_AA_ = 1.7, *m*_AB_ = 1.2, *m*_BA_ = 1.2, *m*_BB_ = 1.7.

Recall that the frequencies are uniform. If we compute directly the information index over the phenotypes *Z*, we get *JZ*_PTI_ = 0.03. The mean propensity for these phenotypes is *M*_z_ = 1.45 and *cov*(*m*_z_, *l*_z_) = 0.0435. However, *M* = 1.5 and *cov*(*m,l*) = 0.1733 for the real mate choice trait (*T*). The scaling is α_z_ = [*cov*(*m*_z_, *l*_z_) /*M*_z_] / [cov (*m, l*) /*M*] = 0.2597 so *J*_PTI_ = *JZ*_PTI_ / α_z_ = 0.03/0.2597 = 0.1155, as expected.

If we have an estimate or a computable proxy for the propensity function m, as for example, a measure of distance between female and male traits |D|, or a model with Gaussian functions (Carvajal-Rodriguez and Rolán-Alvarez, 2014), then *JZ*_PTI_ and *J*_PTI_ can be estimated separately. We obtain *J*_PTI_ by means of *J*(*q*′ *q*) using the estimated mating propensities to classify the frequencies, and we still can use the phenotypes *Z* to compute *JZ*_PTI_. The relationship between both measures may give an idea about the linkage between the phenotypes *Z* and the mate choice.

Suppose that the estimate of *J*_PTI_ is different from zero while *JZ*_PTI_ = 0, then mate choice do exist but it is not linked with the phenotype *Z*. An interested researcher could compare different traits looking for the ones having the best scaling for the information *J*_PTI_, i.e. the one that is more involved in the mate choice. It seems that if we are able of having good proxies for mating propensity, this could pave the way for testing the impact of different traits on mate choice.

Additionally, we still can compute directly Δ*Z* = *Z* – *Z*, i.e. the difference in phenotype frequencies between observed and expected by random mating. Therefore, we have two values, Δ_m_*Z* and Δ*Z*, for the change in *Z*, the discrepancy between them gives an estimate of the change in *Z* caused by other factors than mating propensity (e.g. predators) so *e*_z_ = Δ*Z* – Δ_m_*Z*.

Thus the total change in mean *Z* is

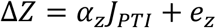

## Appendix B

The relationship between sexual selection measured within sex and the pair sexual selection measured by *PSS* is

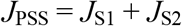

To see this, recall that *J*_PSS_ is the sum of products 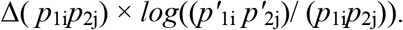. Then note that

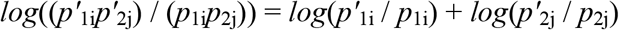

and that

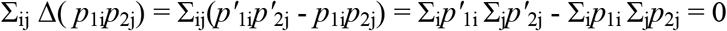

because each summation is 1. Then, after some algebraic rearrangement we obtain

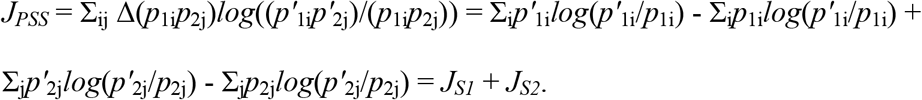

## Appendix C. Scaling factors

We can compute the scaling factors that translate the information between different phenotypic scales.

We have used the notation *J*_PTI_, *J*_PSS_, *J*_PSI_ and *E*_0_ for indicating the information when measured from phenotypes that are the choice targets i.e. the phenotypes that the mates care about in choosing each other. On the other hand, we note *JZ*_PTI_, *JZ*_PSS_, *JZ*_PSI_ and *ZE*_0_ when the phenotypes may or may not be related with the choice.

Therefore, it is interesting to shown how the information changes between one measures or others. So that

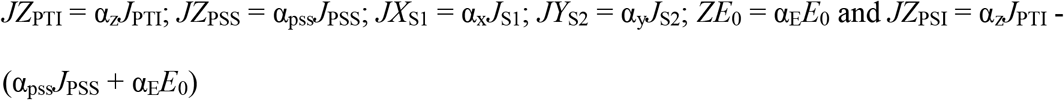

The scalings are as follows.

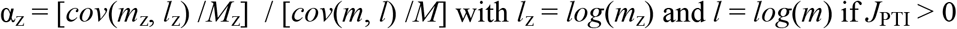

or α_z_ otherwise (*J*_PTI_ = 0)

However, we can also express *cov*(*m*_z_, *l*_z_) /*M*_z_ as β(*m*_z_,*l*_z_)×*D*_z_ which is the regression of the propensity under the trait *Z* over its logarithm multiplied by the index of dispersion. Then if *D*_m_ = *V*(*m*) / *M* and *D*_z_ = *V*(*m*_z_)/*M*_z_ are the indexes of dispersion over the choice and *Z* traits respectively, we obtain

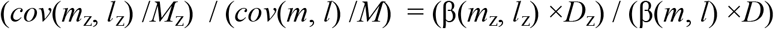

Then if if *J*_PTI_ > 0 define

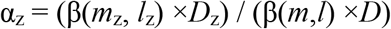

where *l*_z_ = *log*(*m*_z_) and *l* = *log*(*m*)
or α_z_ = 1 if *J*_PSI_ = 0

Similarly,

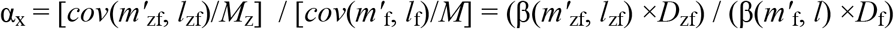

or 1 if *J*_PSI_ = 0
where 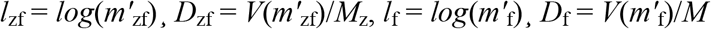

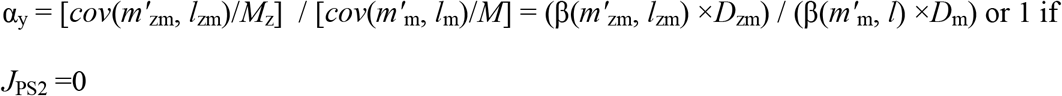

where 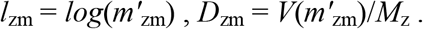

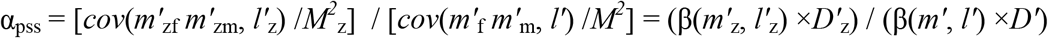

or 1 if *J*_PSI_ = 0
where 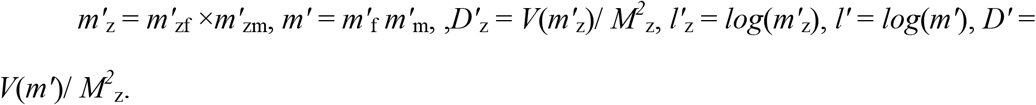

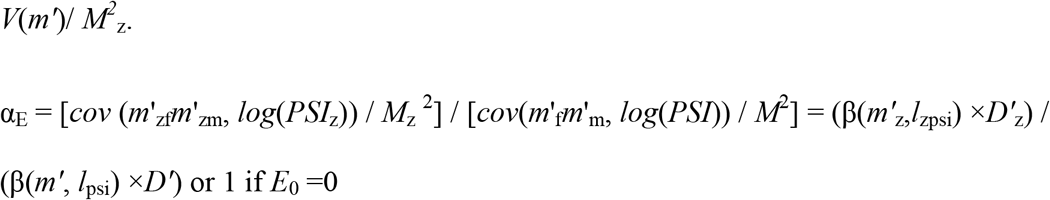

where *l*_zpsi_ = *log(PSI_z_)*, *l*_psi_ = *log*(*PSI*).

Finally

if *J*_PSI_ = 0

α_δ_ = α_z_ when *J*_PSS_ = 0 or in general

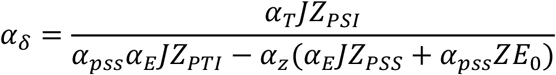

with α_T_ = α_pss_ x α_E_; so that *JZ*_PSI_ = α_β_*J*_PSI_

## Appendix D. Multiplicative preference example

Consider two male traits, *B/b* for acoustic broadcast and *D/d* for display rate, where in both cases the upper case refers to the higher value of the trait; consider also one female trait *X* with a single phenotypic class. The females are the choosers and so we may define multiplicative preferences as appear in Table S1.

**Table S1.**
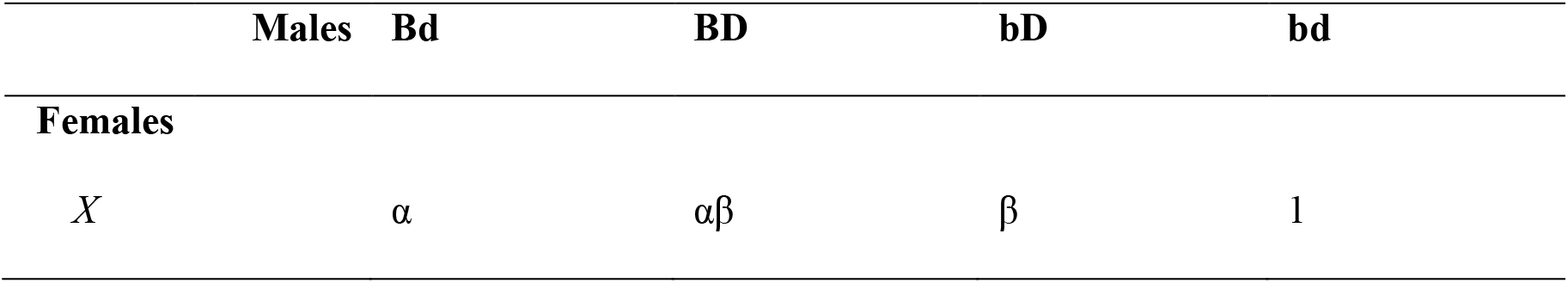
Mating preferences for acoustic broadcast and display rate.

The model assumes a single female phenotypic class and so the frequency is 1. Let the frequencies of the different male classes be *p*_Bd_, *p*_BD_, *p*_bD_ and *p*_bd_. Then the mean preference is *M* = *αp*_Bd_ + αβ*p*_BD_ + β*p*_bD_ + *p*_bd_. There is only one female marginal preference, that in fact coincides with the mean preference, *m*^′^_fX_ = *M*. There are four male marginal preferences 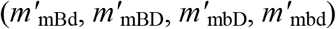 that match the corresponding male propensity with the single female phenotype i.e. 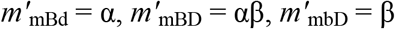 and 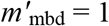.

In (5) we have given an aprioristic expression (δ) of the pair sexual isolation (*PSI*) coefficients in terms of the preferences so that 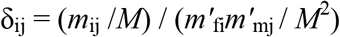. Therefore, 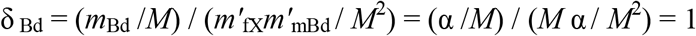 and similarly for the other phenotypes so finally δ_Bd_ = δ_BD_ = δ_bD_ = δ_bd_ = 1, which implies no sexual isolation.

On the other hand, the model predicts male sexual selection whenever α and/or β ≠ 1.

## Appendix E

*Proposition* 1

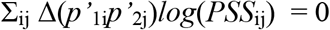

then

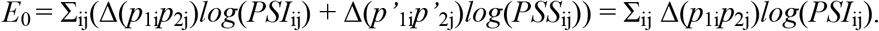

First, recall that

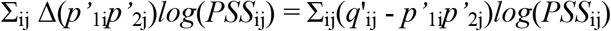

and also that by definition of *PSS*

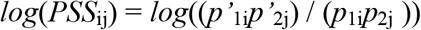

that can be expressed as

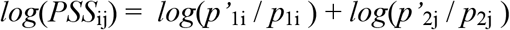

then by simple substitution and rearranging the terms

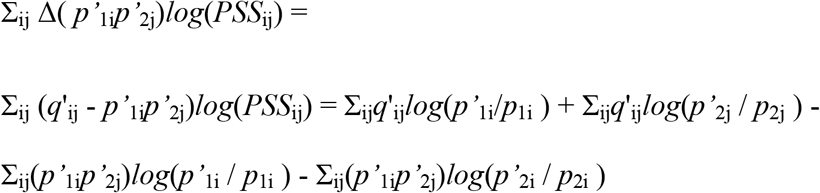

Now recall that the *i* subscript refers to females and subscript *j* to males, then the double summatory is the sum through females and males, thus by reminding that 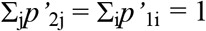 we note that

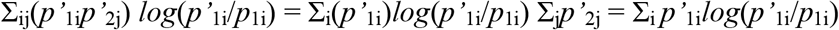

and similarly

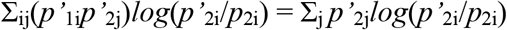

so we have

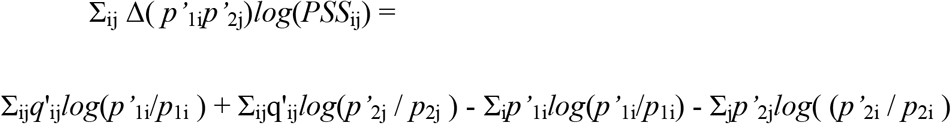

Now note that

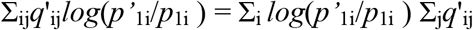

and that for each female *i* the sum through males of the observed mating frequencies involving female *i* is, by definition, *p*′_1i_ i.e. 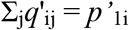 and similarly for each male *j* we have 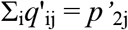. Then

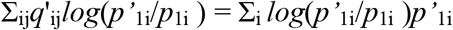

and

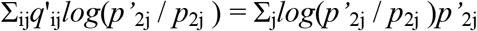

therefore

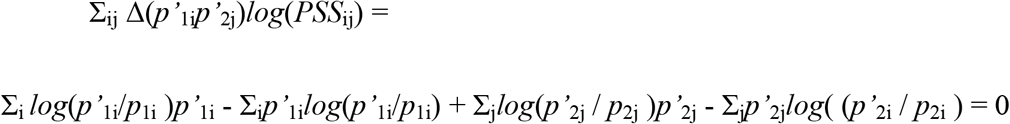

and so the proposition is true

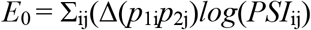

*Proposition* 2

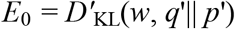

where

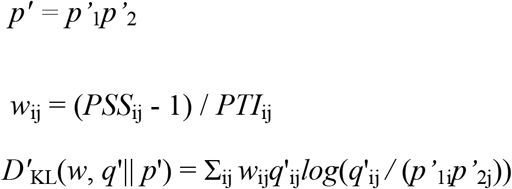

From the model (1) and the partitions (4) and (5) in the main text we know that

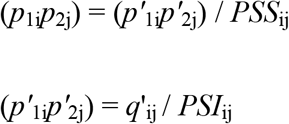

therefore

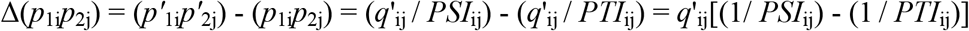

and since *PTI*_ij_ = *PSI*_ij_ x *PSS*_ij_ we obtain

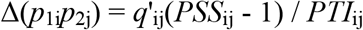

and so

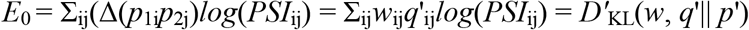

which is Kullback–Leibler-like divergence with weights *w*_ij_ in the observations *q*′.

